# LLM-assisted analytical review catches statistical errors that human researchers miss: documented evidence from two computational biology case studies

**DOI:** 10.64898/2026.10.15.759333

**Authors:** Om Chhaya, Viswanath Ananth

## Abstract

Large language models (LLMs) are increasingly used to assist with code generation and data analysis in computational biology, but their capacity to serve as analytical reviewers — detecting statistical and methodological errors in completed analyses — is less documented. We present two case studies in which LLM-assisted review identified consequential errors in computational biology analyses that the human researchers had not detected. In single-cell transcriptomics, review of a differential expression analysis of 22,048 cells from human striatum identified a sex composition confound (Chi-squared = 192.7, p = 8.1 × 10^−44^) that had produced spurious Y-chromosome gene hits; correction revealed KIF3A as the sole genuine finding (adjusted p = 1.37 × 10^−14^). In HRV-based stress classification, review identified subject leakage in a train-test split that allowed the same individuals in training and test sets; correction via leave-one-subject-out cross-validation produced a more reliable and higher AUC estimate (0.946 vs. 0.848). Both errors would have survived conventional peer review in the absence of systematic checking. Both corrected analyses have been submitted for peer review. These cases suggest that LLM-assisted analytical review — using LLMs to interrogate completed analyses for confounds, leakage, and statistical artifacts — is a practical quality assurance step that complements rather than replaces human scientific judgment.

## Introduction

The use of large language models (LLMs) in computational biology has expanded rapidly, with applications ranging from code generation and literature synthesis to autonomous multi-agent research systems [1,2]. Systems such as BioMedAgent [3], The Virtual Biotech [4], and The AI Scientist [5] demonstrate that LLMs can plan and execute complex analytical workflows. However, much of this work focuses on LLMs as *producers* of analysis. Less attention has been paid to their potential as *reviewers* — systematic checkers of completed analyses for statistical and methodological errors.

This distinction matters because the most consequential errors in computational biology are often not bugs in code but flaws in analytical design: confounds in group comparisons, leakage between training and test sets, inappropriate multiple testing correction, and circular analyses. These errors are difficult to detect because the code runs correctly and produces plausible-looking results. They are caught, when they are caught at all, by experienced reviewers who know which checks to perform — or by peer reviewers after submission, when correction is costly and embarrassing.

We present two case studies in which LLM-assisted review — specifically, using Claude (Anthropic) to interrogate completed analyses — identified consequential errors that the human researchers had not detected. In both cases, the errors would have produced publishable but misleading results. In both cases, the corrected analyses were stronger. We document the errors, the detection process, and the corrected results, and discuss the practical implications for quality assurance in computational biology research.

## Materials and methods

### Analytical review process

In both case studies, the analysis pipelines were developed by the first author (OC) using standard computational biology tools, with code refinement assistance from ChatGPT (OpenAI, GPT-4). After the analyses were complete and results generated, the outputs were reviewed using Claude (Anthropic, claude-opus-4-6) in an interactive conversational format. The review process involved presenting the analytical design, data structure, and results to the LLM and requesting systematic checking for potential confounds, leakage, statistical artifacts, and methodological inconsistencies.

This process is distinct from using LLMs for code generation or pipeline automation. The analyses were already complete. The LLM’s role was to review them — analogous to an internal statistical reviewer or a methods-focused collaborator who examines the analytical design after results are available. All analyses were executed on a standard Windows 10 laptop (Python 3.13, 16 GB RAM, no GPU) with total runtimes under 10 minutes per case study.

### Case study 1: Single-cell transcriptomics

A differential expression analysis was performed on the GSE197140 dataset (22,048 cells from human striatal tissue) to identify genes associated with ciliation status in specific neuronal subtypes. Per-cell ciliation scores were computed using Scanpy’s score_genes function, cells were divided into high and low ciliation groups by within-cell-type median split, and Wilcoxon rank-sum tests with Benjamini-Hochberg correction were applied. The analysis was implemented using Scanpy 1.10 and SciPy 1.13. Full biological results and methods are reported in a companion publication [6].

### Case study 2: HRV stress classification

A heart rate variability (HRV) classification pipeline was developed using the WESAD public dataset (15 subjects, chest-worn ECG under stress, rest, and amusement conditions) [7]. Six HRV features (RMSSD, SDNN, pNN50, MeanNN, LF, HF, LF/HF) were extracted using NeuroKit2 [8], and binary stress classifiers (Random Forest, Logistic Regression, SVM) were trained using scikit-learn [9]. The initial evaluation used a 70/30 stratified train-test split. Full results are reported in a companion publication [10].

## Results

### Case study 1: Sex composition confound in differential expression

The initial analysis produced a ranked list of differentially expressed genes between high and low ciliation groups in neurons. The top three genes were USP9Y, NLGN4Y, and UTY — all located on the Y chromosome.

During LLM-assisted review, the reviewer noted that all three top hits were Y-linked and requested a test for association between the ciliation grouping variable and donor sex. The test revealed a profound confound: Chi-squared = 192.7, p = 8.1 × 10^−44^. The high-ciliation group was heavily enriched for male donors, meaning the ‘ciliation-associated’ gene signature was substantially a sex signature.

The correction involved: (1) excluding 13 Y-chromosome genes from the analysis, (2) repeating the differential expression within each cell type separately, and (3) testing the sex-ciliation association within each cell type. This revealed that PV interneurons were the only cell type where the confound was marginal (p = 0.052), while neurons were heavily confounded (p = 2.3 × 10^−47^). In the corrected PV interneuron analysis, KIF3A emerged as the sole genome-wide significant gene (adjusted p = 1.37 × 10^−14^, log_2_FC = +2.67; Fig 1) — a biologically coherent finding, as KIF3A encodes the essential kinesin motor for primary cilia assembly.

**Fig 1.**
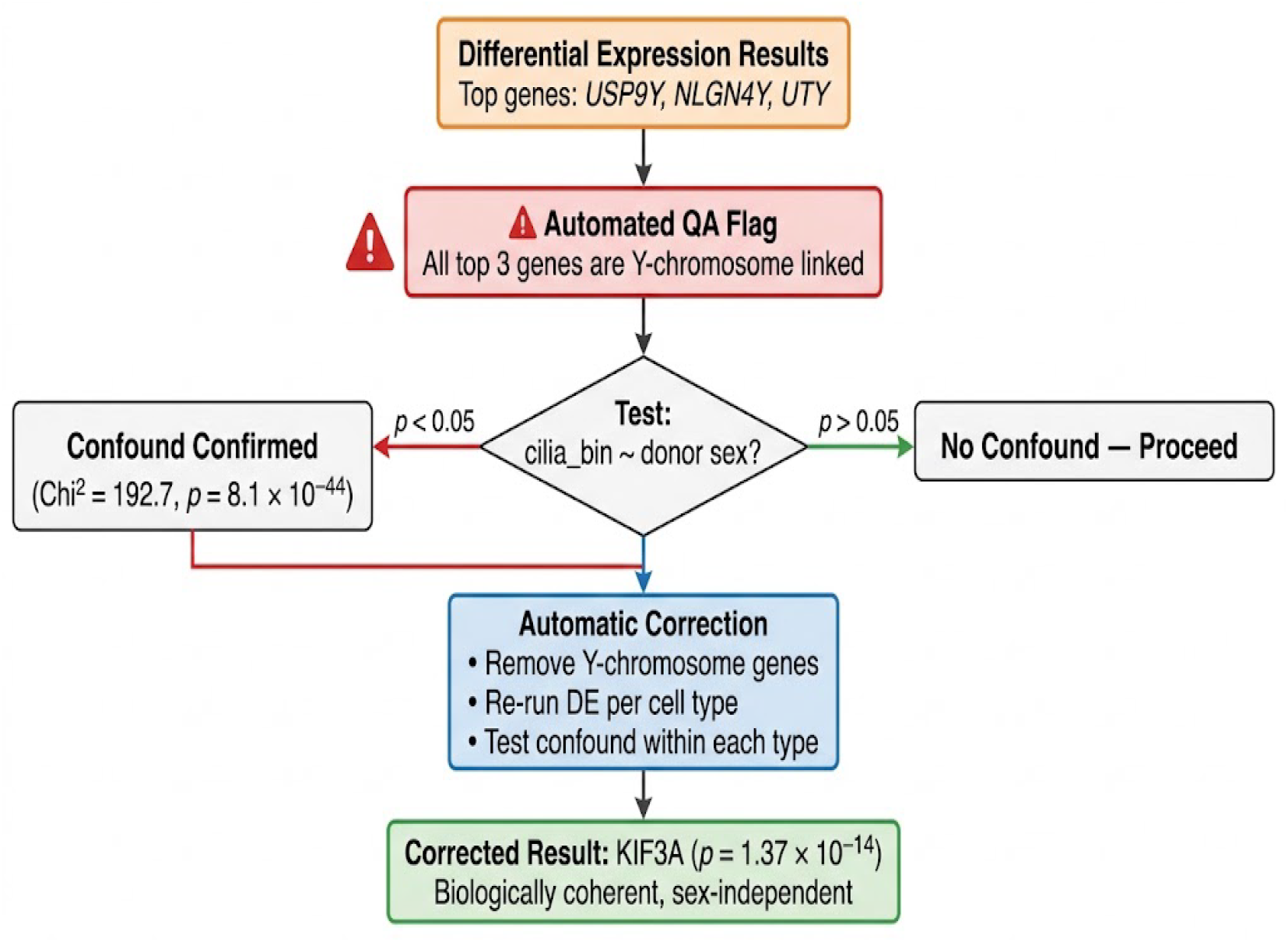
Error detection and correction in Case Study 1. LLM-assisted review identified that all top differentially expressed genes were Y-chromosome linked, triggering a sex-ciliation association test that revealed a profound confound (p = 8.1 × 10^−44^). Correction produced the genuine finding: KIF3A (adjusted p = 1.37 × 10^−14^).

**Fig 2.**
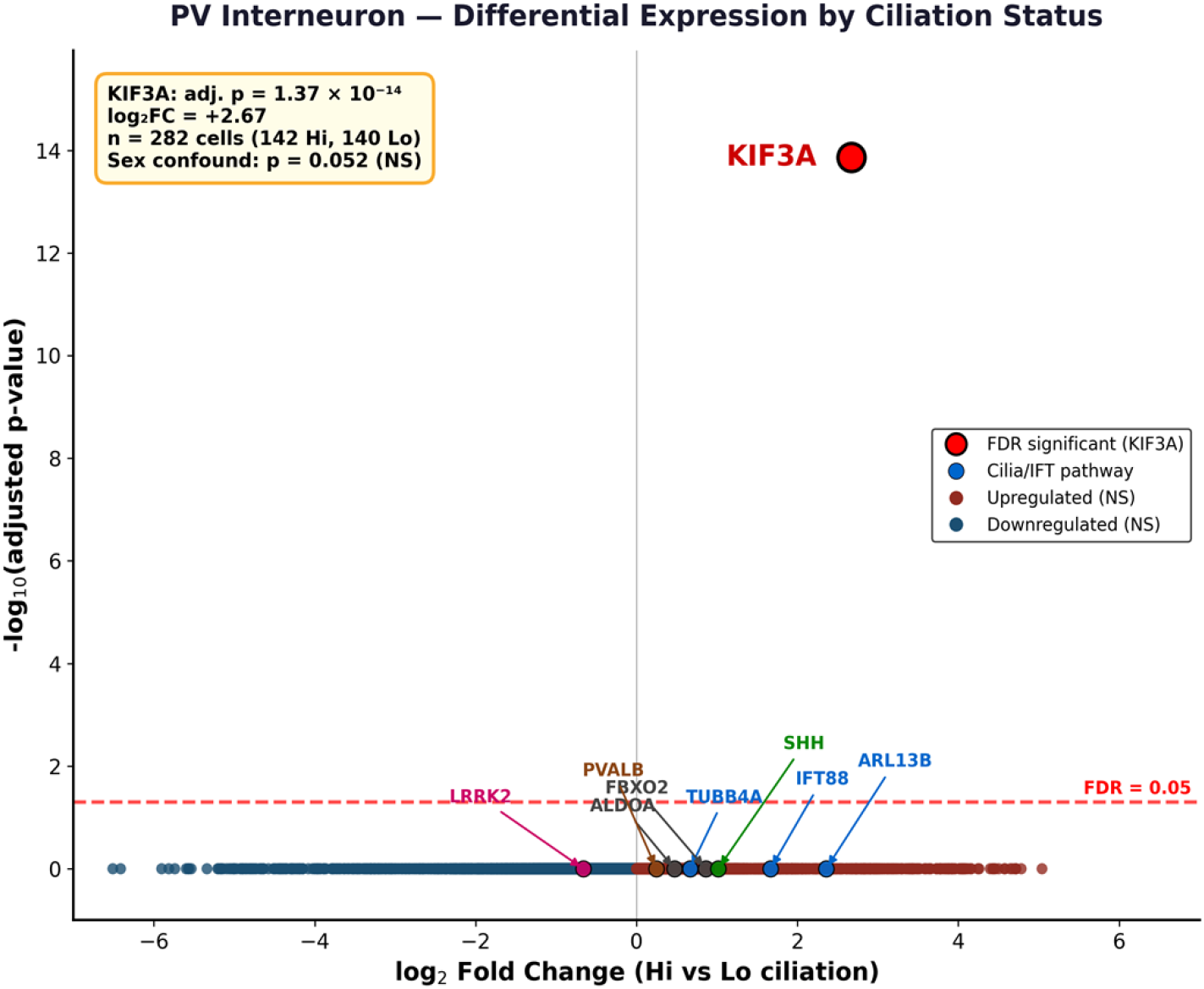
Corrected result: volcano plot of differential expression in PV interneurons. KIF3A is the sole genome-wide significant gene after Y-chromosome exclusion and within-cell-type analysis.

#### Consequence of the error

Without detection, the published analysis would have reported USP9Y, NLGN4Y, and UTY as ‘ciliation-associated’ genes — a spurious finding driven entirely by sex composition. This would not have been obviously wrong to reviewers unfamiliar with the Y-chromosome locations of these genes.

### Case study 2: Subject leakage in train-test split

The initial analysis used a 70/30 stratified train-test split to evaluate HRV classifiers, producing a Random Forest ROC AUC of 0.848. During LLM-assisted review, the reviewer examined the data structure and identified that the 45 observations (15 subjects x 3 conditions) were split by row, not by subject. This meant that the same individual’s rest data could appear in training while their stress data appeared in testing, allowing the classifier to learn person-specific cardiac signatures rather than generalizable stress patterns.

The correction replaced the train-test split with leave-one-subject-out (LOSO) cross-validation across all 15 subjects. Under LOSO, each subject’s data was predicted using a model trained on the other 14 subjects, ensuring no information leakage. The corrected Random Forest AUC was 0.946 (95% CI: 0.871-0.998; Fig 3), substantially higher than the leaked estimate — reflecting a more stable evaluation across all 45 samples rather than a single split of 15 test samples.

**Fig 3.**
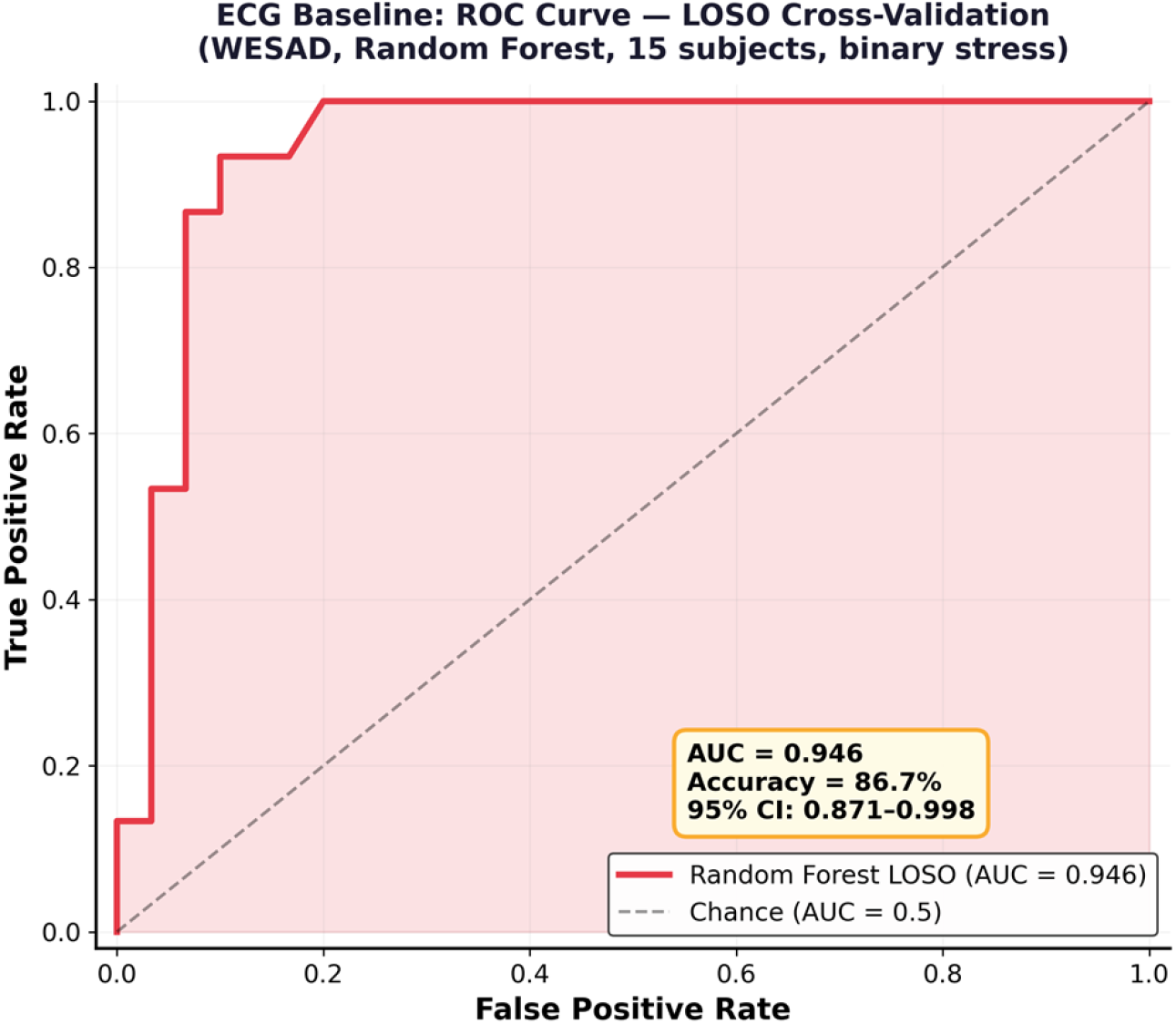
Corrected result: ROC curve under LOSO cross-validation. Random Forest AUC = 0.946 (95% CI: 0.871-0.998). Each of 15 subjects predicted once when held out.

**Fig 5.**
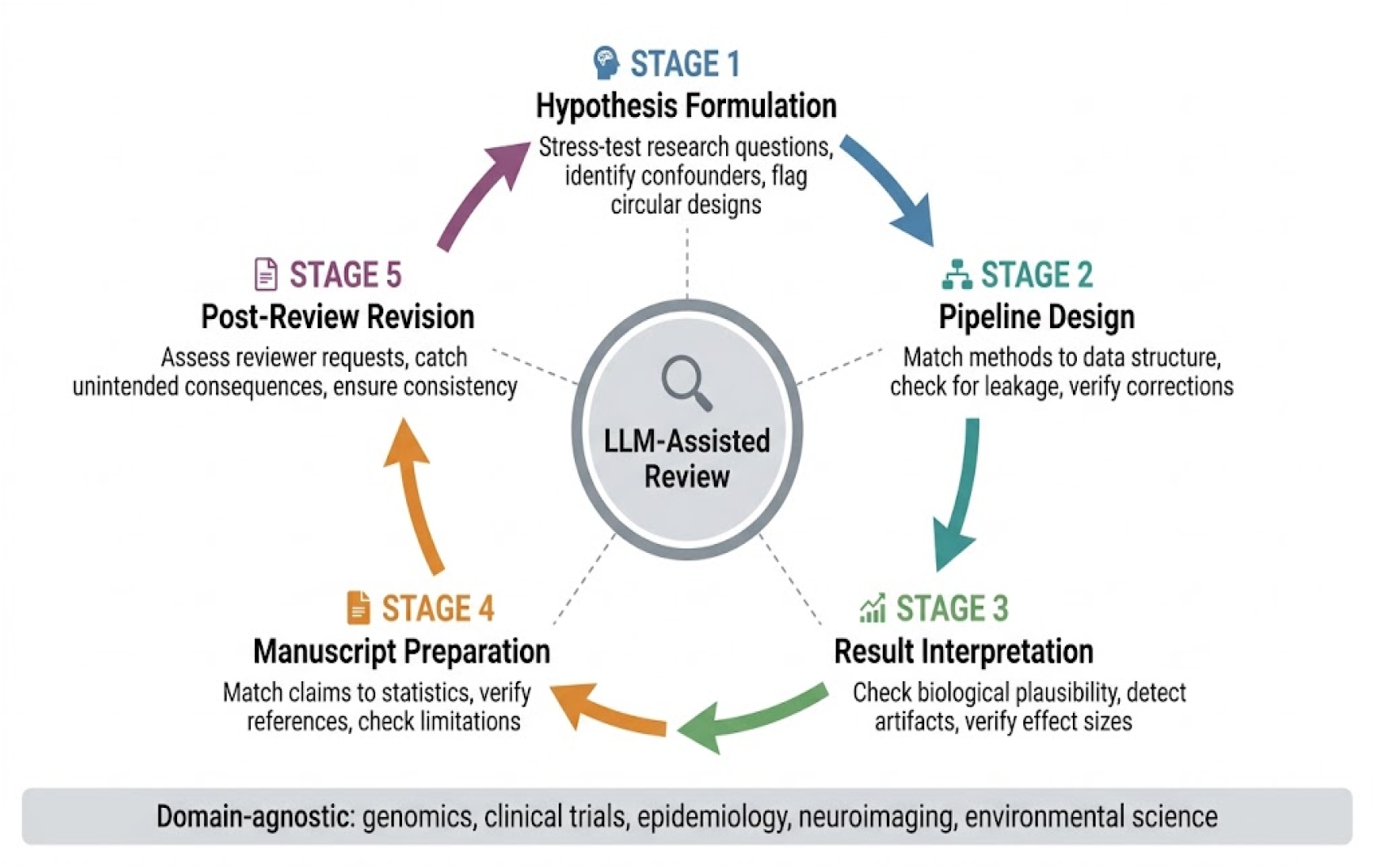
LLM-assisted review as a continuous feedback loop across the research workflow. Five stages from hypothesis formulation through post-review revision, each connected to a central LLM review hub. The approach is domain-agnostic and iterative — corrections at any stage can trigger re-evaluation at earlier stages.

#### Consequence of the error

The leaked AUC of 0.848, while lower than the corrected 0.946, was not inflated in the usual sense. However, it was unreliable — a single random split of 15 test samples producing a point estimate with high variance. A different random seed would have produced a substantially different number. The LOSO estimate is both more reliable and more interpretable, as it directly answers the clinical question: does the classifier generalize to individuals it has never seen?

### Summary of errors detected

**Table 1.**
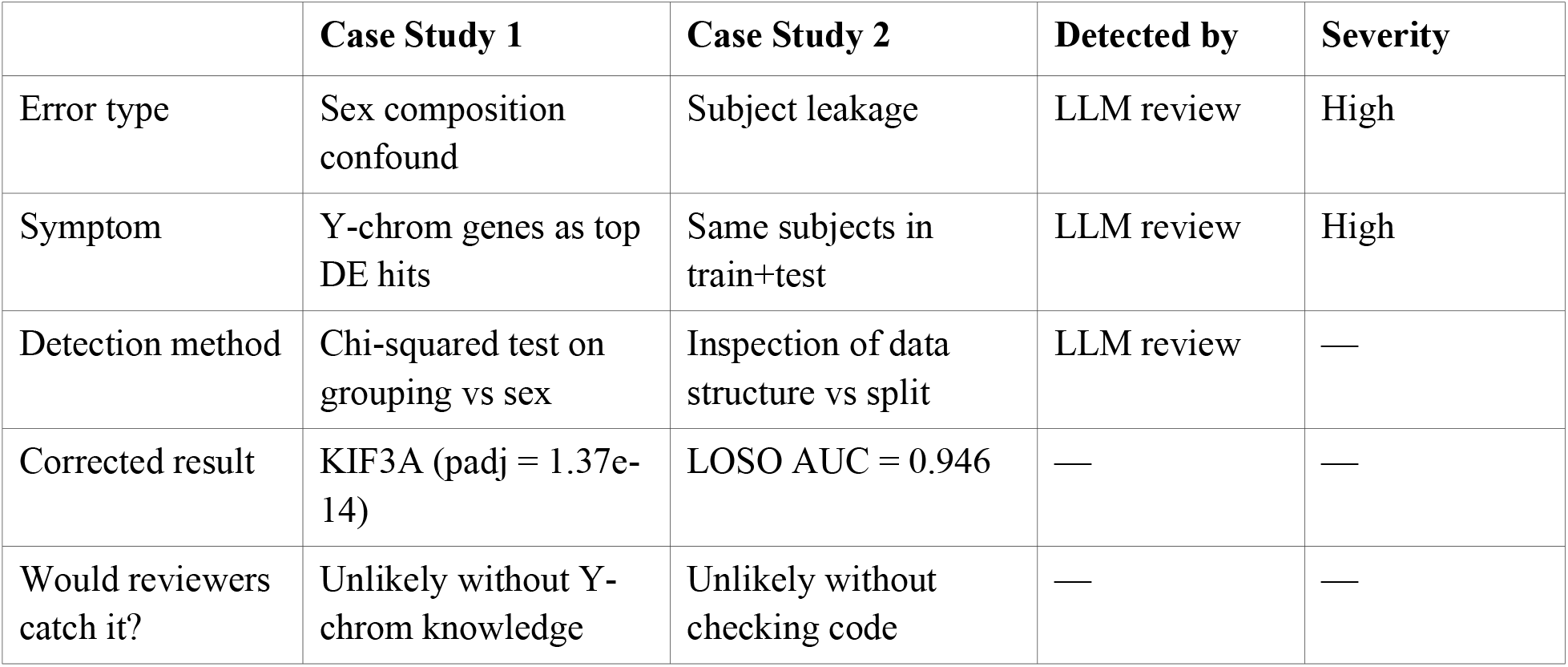
Summary of errors detected by LLM-assisted analytical review.

## Discussion

These two case studies illustrate a practical role for LLMs in computational biology that is distinct from code generation, pipeline automation, or autonomous research: systematic analytical review of completed work. In both cases, the human researcher had completed the analysis, generated results, and was preparing for submission. In both cases, the LLM reviewer identified a consequential error by performing a check that the researcher had not thought to perform.

The sex confound in Case Study 1 is a textbook example of Simpson’s paradox — an association driven by a lurking variable. The researcher did not test for it because the analysis appeared to produce biologically plausible results (ciliary-related genes). The LLM reviewer flagged it because all three top genes happened to be Y-linked, which triggered a systematic check. This is the kind of pattern recognition that experienced bioinformaticians perform routinely but that less experienced researchers may miss.

The subject leakage in Case Study 2 is one of the most common errors in biomedical machine learning [11]. It produces results that look correct — the code runs, the metrics are reasonable, the figures are clean — but the evaluation is invalid because the model has implicitly seen the test subjects during training. The LLM reviewer caught it by examining the data structure (15 subjects, 3 conditions each, 45 rows) against the splitting strategy (row-level stratified split). This is a mechanical check that could be automated, but in practice is frequently missed.

### What LLM-assisted review is and is not

We are not proposing LLMs as autonomous quality assurance systems. The review process in both case studies was interactive and conversational — the human researcher presented the work, the LLM asked questions and suggested checks, and the researcher executed the corrections. The LLM did not run code, access data, or make scientific decisions. It served as a systematic, tireless interlocutor who knew which questions to ask.

This role is closest to that of a statistical reviewer or a methods-focused collaborator — someone who examines the analytical design with fresh eyes and domain knowledge of common pitfalls. The practical advantage of an LLM in this role is availability: it does not require scheduling, it does not have competing priorities, and it can review work at any hour. The limitation is that it cannot verify claims against the actual data — it reviews the logic and design of the analysis as described, not the data itself.

### Limitations

Several important limitations apply. First, two case studies do not establish the generalizability of LLM-assisted review. The errors detected here — sex confounds and subject leakage — are well-known pitfalls that any experienced reviewer might catch. Whether LLMs can detect subtler or more domain-specific errors remains untested. Second, the review process depends on the human researcher accurately describing their analytical design; if the description is incomplete or misleading, the LLM cannot catch errors it cannot see. Third, LLM outputs are non-deterministic — a different conversation might not have surfaced the same checks. Fourth, we tested only one LLM (Claude, Anthropic); generalization across models is not established, though the code refinement used a different LLM (ChatGPT, OpenAI), suggesting the approach is not model-specific. Fifth, both companion publications are under peer review; acceptance is not guaranteed.

### Practical recommendations

Based on these experiences, we suggest that computational biology researchers consider adding an LLM-assisted review step before submission. The process requires no specialized tools — only an interactive conversation with a frontier LLM in which the researcher describes the data structure, analytical design, grouping variables, splitting strategy, and top results, and asks the LLM to identify potential confounds, leakage, circularity, and statistical artifacts. This step takes approximately 30-60 minutes and complements — but does not replace — internal review by collaborators and formal peer review.

### Broader applicability across the research workflow

While the case studies presented here focus on post-analysis review — catching errors after the pipeline has run — the same LLM-assisted approach applies across the entire research workflow as a continuous feedback loop. We outline five stages where interactive LLM review adds value:

#### Hypothesis formulation

Before any code is written, an LLM can stress-test a research question against existing literature, identify potential confounders inherent in the proposed study design, and flag circular or unfalsifiable hypotheses. In Case Study 1, a pre-analysis review might have identified the sex confound risk in the GSE197140 dataset before the first line of code was executed.

#### Analytical pipeline design

When building the data processing and statistical pipeline, LLM review can evaluate whether the chosen methods match the data structure — for example, whether a classification evaluation strategy accounts for grouped observations (as in Case Study 2), whether multiple testing correction is appropriate for the number of comparisons, or whether feature engineering introduces circularity. This is the stage where the most consequential design decisions are made and where errors are cheapest to fix.

#### Result interpretation

After results are generated, LLM review can examine whether top findings are biologically plausible, whether effect sizes are consistent with the sample size, whether confidence intervals are appropriately constructed, and whether the results could be explained by artifacts. Both case studies demonstrate this stage: the Y-chromosome pattern in Case Study 1 and the train-test structure in Case Study 2 were visible in the results but required systematic checking to detect.

#### Manuscript preparation

Before submission, LLM review can check whether claims in the text match the reported statistics, whether figure captions accurately describe what is shown, whether references support the sentences that cite them, and whether limitations are honestly stated. This is the stage closest to traditional peer review, but performed before the manuscript leaves the authors’ hands.

#### Post-review revision

When reviewer feedback arrives, LLM-assisted review can help assess whether requested changes are scientifically sound, identify unintended consequences of revisions (such as a corrected analysis that invalidates a downstream claim), and ensure consistency across the revised manuscript.

Crucially, these stages are not sequential but iterative. The correction of subject leakage in Case Study 2 required not only re-running the classification (pipeline design) but also reinterpreting the results (result interpretation), rewriting the manuscript (manuscript preparation), and updating the companion methodology paper (post-review revision). An LLM-assisted review loop that operates across all five stages — from hypothesis through publication — provides continuous quality assurance that catches errors at the earliest and least costly point in the workflow.

This workflow-spanning approach is domain-agnostic. The same LLM-assisted review process applies to genomics, proteomics, clinical trials, epidemiology, neuroimaging, environmental science, and any field where computational analyses mediate between raw data and published claims. The requirement is not domain-specific tooling but a structured conversation between a researcher who understands the science and an LLM that knows which analytical checks to suggest.

## Conclusions

LLM-assisted analytical review detected a sex composition confound and a subject leakage error in two computational biology analyses that had been completed and prepared for submission. Both errors would have produced publishable but misleading results. Both corrections strengthened the analyses. Beyond post-analysis error detection, the same interactive review approach applies across the entire research workflow — from hypothesis formulation through pipeline design, result interpretation, manuscript preparation, and post-review revision — providing a continuous, iterative quality assurance loop. The approach is domain-agnostic, requires no specialized infrastructure beyond access to a frontier LLM, and complements rather than replaces human scientific judgment.

## Use of generative AI

This paper describes the use of LLMs as analytical reviewers. Analysis pipelines were developed by the first author with code refinement from ChatGPT (OpenAI, GPT-4). Analytical review was performed using Claude (Anthropic, claude-opus-4-6) in interactive conversation. All scientific decisions and corrections were executed by the authors.

## Data availability

Case Study 1: GSE197140 (GEO), analysis code at https://github.com/Om-C2026/lrrk2-kif3a-ciliation-pv-interneurons, processed data at doi:10.5281/zenodo.23010404. Case Study 2: WESAD (https://ubicomp.eti.uni-siegen.de/home/datasets/icmi18/), analysis code at https://github.com/Om-C2026/hrv-aud-calibration.

## Acknowledgments

The authors thank Prof. Scott Pappada (Department of Bioengineering, University of Toledo) for mentoring and access to wearable devices used in Case Study 2.

